# Injection of antibodies against immunodominant epitopes tunes germinal centres to generate broadly neutralizing antibodies

**DOI:** 10.1101/501247

**Authors:** Michael Meyer-Hermann

## Abstract

The development of broadly neutralizing antibodies is critical for the control of many life-threatening viral infections like HIV, influenza, or hepatitis. Elite neutralizers are generated by natural germinal centre (GC) reactions in a very small subset of patients. The best strategy of how to promote the generation of broadly neutralizing antibodies is not known. Here, computer simulations of GC reactions are used to predict a feedback loop based on memory B cells from previous GC reactions that promotes GCs to focus on new epitopes. The simulations suggest that the injection of antibodies against immunodominant epitopes fosters affinity maturation of antibodies specific for rare or hidden epitopes. This principle can be used for the design and testing of future therapies and vaccination protocols.

## Introduction

In vaccine research, major efforts aim at inducing the generation of broadly neutralizing antibodies (bnAbs) against viral epitopes. BnAbs are either cross-reactive against different epitopes in the hypervariable region or target a conserved region of the viral antigen (Mascola et al., 2013). They occasionally develop in patients and target different epitopes of the viral antigen (Scheid et al., 2009). It was shown for HIV that one may design bnAbs in mice by sequential immunisation with slightly modified antigen (Escolano et al., 2016). In this way, a sequence of multiple germinal centre (GC) reactions is induced which targets an antigen-epitope converging to the desired bnAb in small steps. There is currently no strategy of how to foster the generation of bnAbs in single GC reactions and vaccination research so far failed to induce bnAbs by immunisation (Schiffner et al., 2013). For example, vaccination against Influenza is often not successful, because the antibodies target highly variable epitopes on the virus surface molecule hemagglutinin (Smith et al., 2004), which is only effective when the viral strain matches the targets (Tricco et al., 2013).

In order to avoid antigenic drift and viral escape it may be more advantageous to develop antibodies targeting the conserved regions of the respective virus. Conserved regions are essential for virus functionality such that viral escape by mutations in these regions would induce a major inhibition in viral entry or replication. Indeed, in the case of HIV, bnAbs target epitopes in the conserved region of the antigen (Kwong et al., 2018). However, these conserved targets are difficult to access by antibodies because the variable regions act as a shield and hide the conserved regions. In GCs, the native antigen is presented on follicular dendritic cells (FDCs) such that in principle it might be possible to find affinity maturation for conserved epitopes, which explains that bnAbs are occasionally found in humans. However, shielding by the immunodominant variable region makes interactions of B cell receptors with conserved epitopes rare and prevents the development of high affinity antibodies in most cases.

There is no existing strategy of how to tune GC reactions to focus on one or the other epitope. Mathematical modelling and computer simulations have been employed before to analyse the generation of high affinity and cross-reactive antibodies (de Boer & Perelson, 2017, Meyer-Hermann et al., 2012, Oprea & Perelson, 1997, Wang et al., 2015). Mathematical models validated by a large set of experimental data are suitable instruments to predict possible ways of modulating the focus of GC reactions. In an elegant study, it was found that pre-existing antibodies might prevent the generation of bnAbs (Zarnitsyna et al., 2016). Indeed, GC-derived or injected antibodies enter the GC reaction and bind to the antigen presented on FDCs (Zhang et al., 2013). They compete with the B cell receptor for binding the target epitope and prevent B cell selection in a concentration- and affinity-dependent way. Even though the presence of immune complexes on FDCs seems dispensable for affinity maturation (Hannum et al., 2000), the presence of high-affinity antibodies can strongly inhibit or shutdown the GC reaction (Zhang et al., 2013). Here, the impact of antibodies specific for immunodominant viral epitopes on affinity maturation of antibodies specific for hardly accessible epitopes is investigated with computer simulations.

## Methods

A GC simulation platform was developed that enables the analysis of affinity maturation in the presence of multiple antigen epitopes. Starting from a state-of-the-art GC simulation software (Meyer-Hermann et al., 2012, Meyer-Hermann, 2014, Papa et al., 2017), in which cell objects are defined in a three-dimensional space, antigen epitopes and antibodies were represented in an abstract affinity space (Perelson & Oster, 1979) heuristically reflecting physical binding properties of antibody and antigen. Cells can move, interact, differentiate, divide, mutate, and die according to generally accepted knowledge about mechanisms of B cell evolution (Victora & Nussenzweig, 2012). Cellular objects include FDCs, dividing B cells, B cells in the state of apoptosis versus selection, and T follicular helper cells (Tfhs). B cells can sensitize and desensitize for chemokines specific for both GC zones and cell motility and the frequency of migration between the zones in the model (Figge et al., 2008, Binder & Meyer-Hermann, 2016) is in accordance with two-photon experiments (Allen et al., 2007, Hauser et al., 2007, Schwickert et al., 2007, Victora et al., 2010).

B cells need to collect antigen from multiple contacts with FDCs for survival. Each antigen epitope is presented on each FDC site at different fractions in order to distinguish high and low accessible epitopes. It is assumed that the probability for a B cell to bind an epitope depends on the affinity of the B cell receptor for the epitope and the accessibility of the epitope. Thereby, the B cells attempt to bind the epitope of highest affinity (default) or of highest accessibility. If the latter option was used, this is explicitly mentioned. Optionally, antibodies, either injected or generated by GC-derived plasma cells, both compete with B cells for binding the epitopes following classical chemical kinetics equations. The antibodies reduce the accessibility of the respective epitopes and by this reduce the probability for a B cell to bind the antigen.

Selection of B cells for re-entering cell cycle or final differentiation to plasma- or memory-cells requires Tfh signalling (Meyer-Hermann et al., 2006, Victora et al., 2010). A threshold Tfh signal has to be achieved by integration of signals from multiple T-B-interactions of 6 minutes each (Papa et al., 2017, Wang et al., 2016). Tfh signalling intensity increases with the amount of collected antigen, i.e. with the density of pMHC presentation by B cells. Selected B cells return to the dark zone for further division and mutation, where the number of divisions depends on the amount of collected antigen (Gitlin et al., 2014, Meyer-Hermann et al., 2012). For a more detailed description of the GC simulation, please refer to the Supplementary Material.

## Results

Some characteristics of the reference simulation are summarised in Figure S1. With a single antigen-epitope in the GC reaction, one can find a diversification of B cell receptors at day three, which is then confined to higher affinity to the presented epitope over the time of the reaction (Figure S1D-F). In the presence of a second related epitope with the same frequency, the GC gives rise to B cells specific to each epitope (Figure 1A). Thus, a single GC environment can give rise to two widely independent affinity maturation processes to two different epitopes.

**Figure 1:**
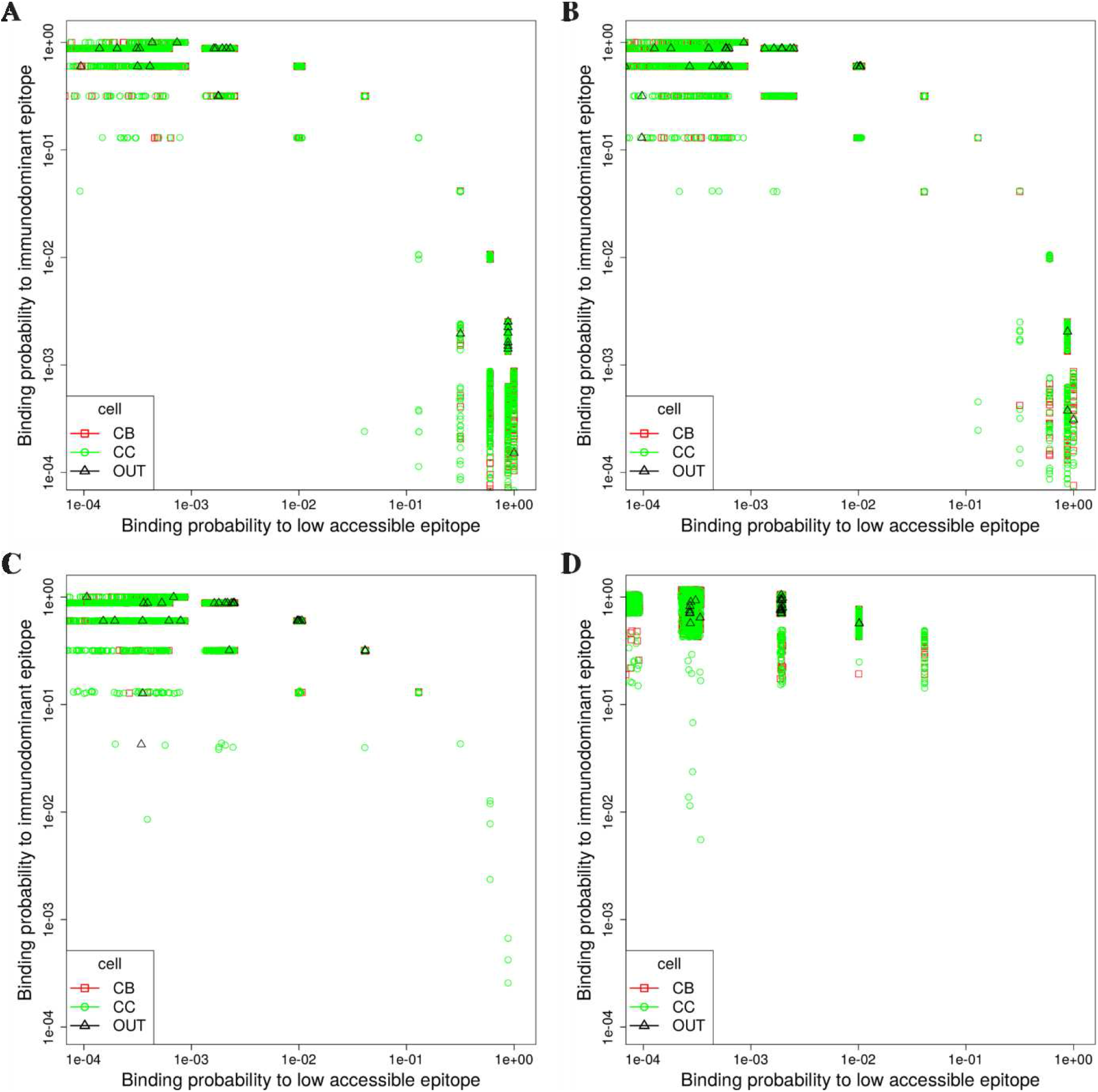
Affinity maturation requires a critical accessibility of the target epitope. *in silico* GC affinity maturation at day 11 post GC onset under the presence of two epitopes with different levels of relative accessibility: 50:50 (A), 30:70 (B), 10:90 (C), 3:97 (D). Both epitopes are unrelated to each other (Hamming distance of 8 mutations). The colour code distinguishes GC-B cells (dividing cells: red open squares; cells in selection mode: green open circles) and GC output cells (black open triangles).

A hardly accessible epitope in the conserved region is mimicked *in silico* by a reduced frequency of its presentation on FDC sites. In the simulations, GC affinity maturation concentrates on the immunodominant epitope (Figure 1B-D). When the probability of catching the low accessible epitope drops to 10% or lower, no affinity maturation against this epitope is observed (Figure 1C,D). This shows that under the presence of an immunodominant epitope, affinity maturation to other epitopes requires a critical minimum accessibility.

The *in silico* GCs in Figure 1 ignore the antibodies generated by the GC reaction output. However, it was shown before that the product of the GC reaction, antibody forming plasma cells, not only efficiently produce antibodies but that these antibodies distribute over the whole organism. In particular, they enter the GC reactions from which they are derived and compete with GC B cells for binding antigen on FDCs (Zhang et al., 2013). In order to analyse the effect of these competitive antibodies, it is assumed that the antibody concentration found in each *in silico* GC stems from the sum of the output cells of all GC reactions in the organism and is diluted over the whole blood and lymph system. A corresponding reference simulation is characterised in Figure S2. At the critical accessibility ratio of 10:90 (Figure 1), the feedback of antibodies induces affinity maturation for the low accessible epitope (Figure 2A). This happens in a late phase of the GC reactions when affinity maturation against the immunodominant epitope is completed and sufficient specific antibody was generated to mask the immunodominant epitope in the GC and increase the likelihood of binding the second epitope. However, the amount of generated output cells is insufficient for a protective immune response. Continuation of the GC simulation for another 3 weeks reveals that the GC response against the low accessible epitope does not take off again (data not shown). These results show that the presence of antibodies specific for the immunodominant epitopes influences affinity maturation for low accessible epitopes but that the naturally produced antibodies hardly induce a full-scale response against those.

**Figure 2:**
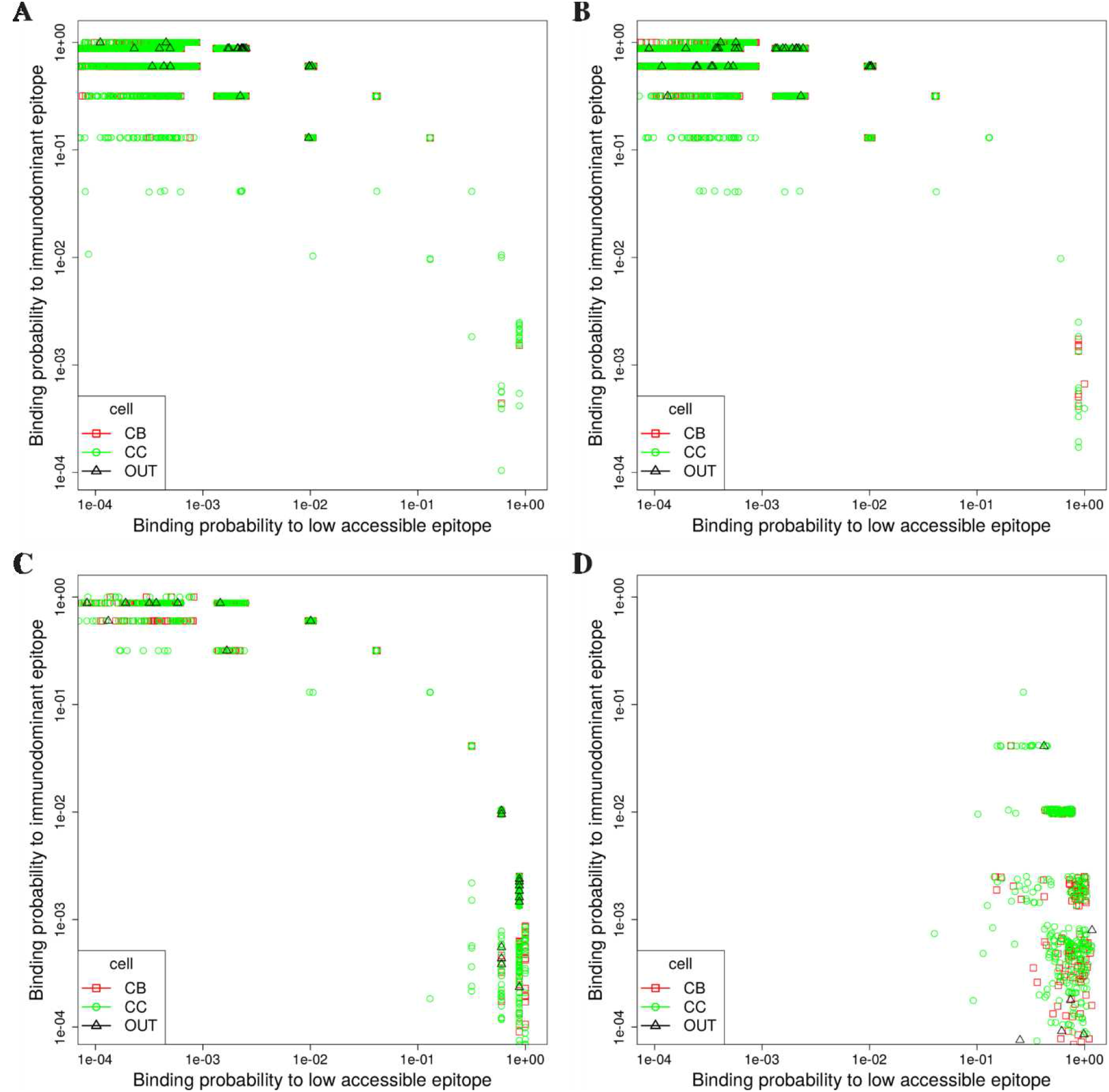
Affinity maturation to two epitopes under the presence of specific antibodies. *in silico* GC affinity maturation to two epitopes with 10:90 accessibility at day 11 post GC onset under the presence of specific antibodies against the immunodominant epitope (vertical axis). (A) Antibodies are derived from the GC reaction itself; (B)-(D) Antibodies are injected at day 3 post GC onset with concentrations of 0.5, 5, 50 nMol, respectively. Both epitopes are unrelated to each other (Hamming distance of 8 mutations). The colour code distinguishes GC-B cells (green open circles) and GC output cells (black open triangles).

The feedback of antibodies generated by the GC output itself might be too late to shift the focus of the GC reaction to a different epitope. Injection of high affinity antibodies specific for the immunodominant epitope at the beginning of the *in silico* GC reaction can stabilize the generation of high affinity output cells specific for the low accessible epitope (Figure 2B-D, Table 1). This effect is also reflected in the total antibody generated against the low accessible epitope (Figure 3) and is dependent on the concentration of the injected antibody against the immunodominant epitope. For low concentrations, the injected antibody increases the relative accessibility of the low accessible epitope. Affinity maturation against both epitopes co-exist. At intermediate antibody concentrations, the affinity maturation for the low accessible epitope becomes dominant. When the generated antibodies cover the anyway low accessible epitope, a second wave of affinity maturation against the immunodominant epitope is deliberated. At high concentrations of injected antibody, affinity maturation for the immunodominant epitope is fully suppressed. Because of the low accessibility, the total size of the GC response is smaller compared to a GC reaction against the immunodominant epitope. However, the probability of selection of B cells specific for even lower accessible epitopes (3%) is consistently increased by the injection of antibodies (Table 1). The result that the injection of antibodies specific for the immunodominant epitope can shift GC affinity maturation to low accessible epitopes turned out robust against model assumptions on antigen uptake (Figure S3) and GC-B cell exit (Figure S4).

**Table 1:** Memory B cell-derived self-inhibition of GC reactions. *in silico* GC simulations with a ratio of immunodominant to low accessible epitope of 97:3. Different settings are compared: **Ab-feedback** is the reference simulation in Figure 2 and Supplementary Figure S2; **Ab-injection** correspond to two settings in Figure 3 with injections of different doses of antibodies specific for the immunodominant epitope; **memory cells** assumes the presence of memory B cells from a previous GC reaction, where these constitute a fraction of the GC founder cells (mdGC) and contribute to early memory-derived antibody-forming plasma cells (mdPC). The number of mdPC should be interpreted as a strength of antibody-production rather than as a number of cells. The readouts comprise the frequency of GCs in which B cells with highest affinity (dissociation constant *K*_D_ < 0.3 nMol) for the low accessible epitope **occurred**. These would be a result of mutation and are counted irrespective of their fate and their frequency in the GC; the frequency of GCs in which B cells specific for the low accessible epitope occurred and were **selected** at least once; and the percentage of GCs classified as *selected* among those classified as *occurred*. 960 simulation for each condition.

| setting/readout | Ab-feedback | Ab-injection |  | memory cells |  |  |  |
| --- | --- | --- | --- | --- | --- | --- | --- |
| number of GCs | 960 | 960 | 960 | 960 | 960 | 960 | 960 |
| memory GC founder | 0 | 0 | 0 | 90% | 25% | 5% | 5% |
| mdPC | 0 | 0 | 0 | 1000 | 1000 | 1000 | 10000 |
| Ab-injection [nMol] | 0 | 5 | 50 | 0 | 0 | 0 | 0 |
| occurred in | 199 | 321 | 248 | 22 | 129 | 200 | 331 |
| selected in | 72 | 168 | 136 | 2 | 24 | 58 | 184 |
| %selected | 36 | 52 | 55 | 9 | 19 | 29 | 56 |

**Figure 3:**
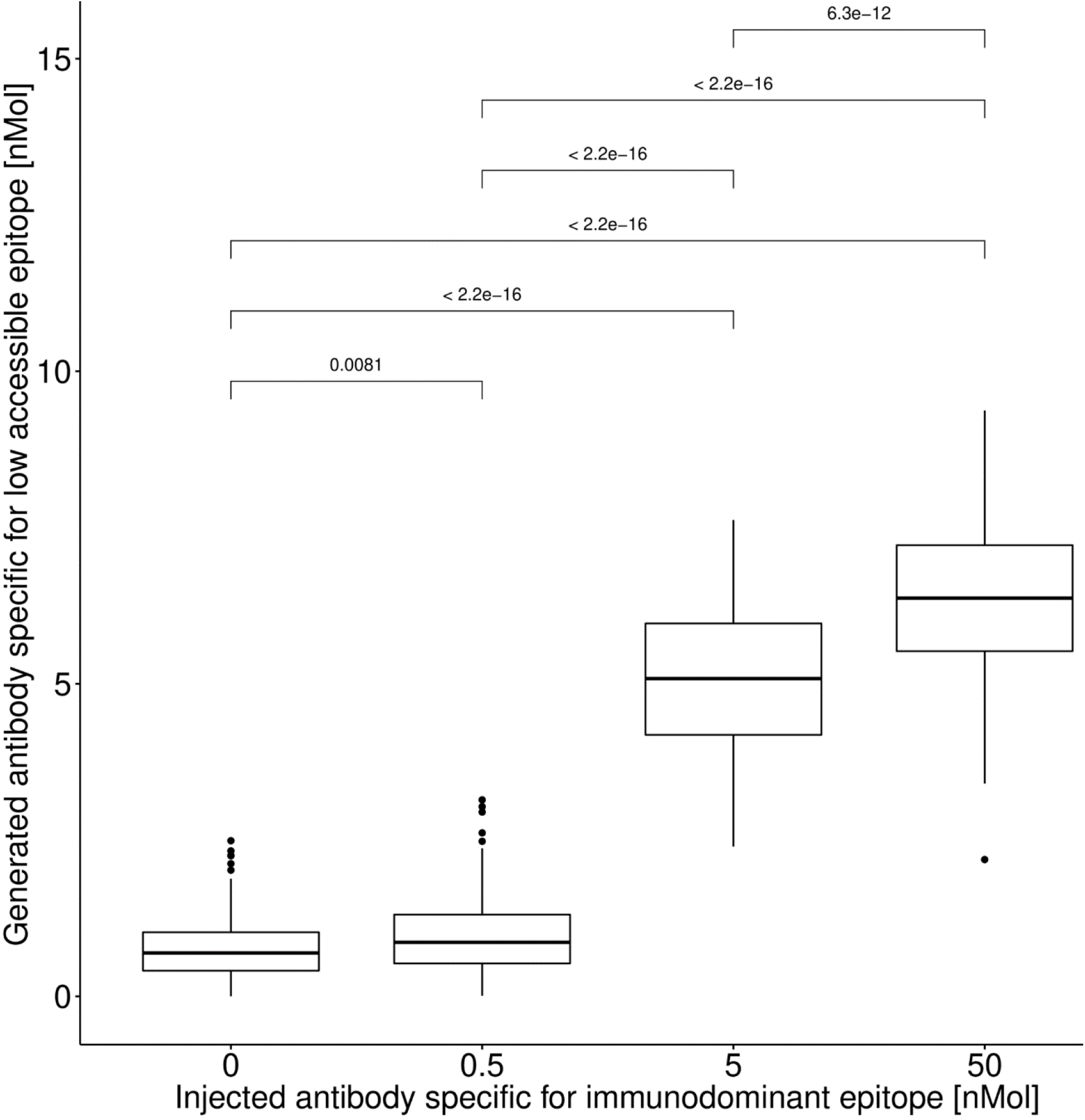
Antibody injection induces more antibodies against low accessible epitopes. The total amount of antibodies specific (dissociation constant *K*_D_ < 2 nMol) for the low accessible epitope generated by the single *in silico* GC reaction at day 21 post GC onset (i.e. at the end of the reaction) without (left) and with different concentrations (0.5, 5, 50 nMol) of injected antibodies specific for the immunodominant epitope. Boxplots of 120 simulations for each condition.

The GC reaction also generates memory B cells. At the time of initiation of a subsequent GC reaction, memory BCs would either differentiate to antibody forming plasma cells or participate in the GC reaction. The B cell receptors of the memory B cells in the GC face highly competent antibodies stemming from the memory derived plasma cells, which compete for antigen binding. The model predicts that memory B cells might well participate in GC reactions but will have a competitive disadvantage as compared to unrelated naïve B cells (Figure 4). Even though both target epitopes are presented at equal frequencies in these simulations, memory B cells are removed from the GC reaction earlier than new B cells unless they managed to survive until they acquired a high number of mutations. The strength of this self-suppression depends on the fraction of memory B cells differentiating to antibody-forming cells (Figure 4). Thus, antibody-feedback provides an additional mechanism of immune memory that allows to progress antibody responses against various epitopes rather than to become trapped in repetitive GC reactions against the same epitope. This mechanism is likely responsible for the occasional natural development of bnAbs.

**Figure 4:**
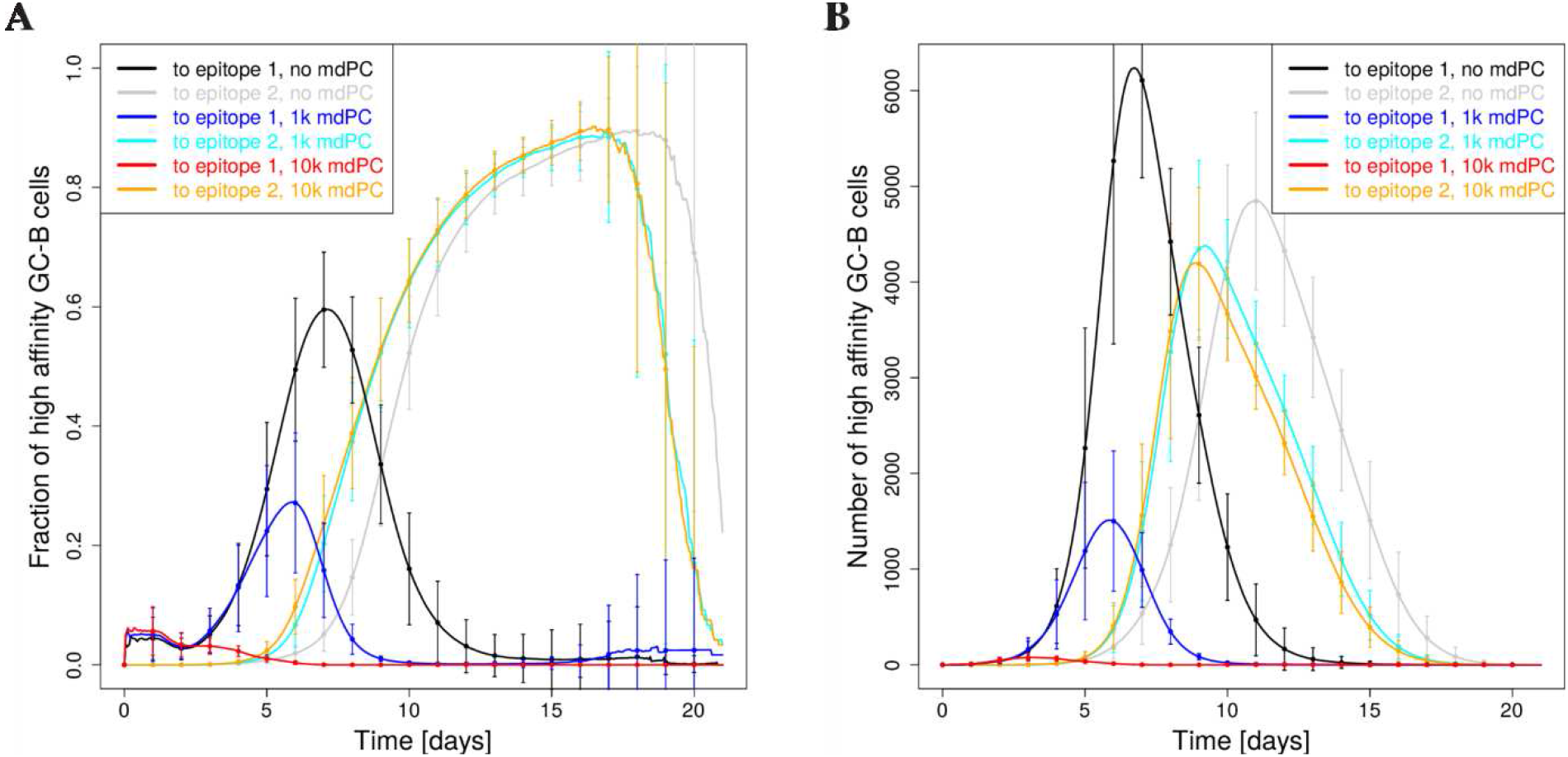
Impact of memory BCs on secondary GC reactions. Fraction (A) and number (B) of high affinity GC-BCs with a dissociation constant of *K*_D_ < 2 nMol. The simulations start with 5% founder cells from a memory BC pool specific for epitope 1. Epitope 2 (at Hamming distance of 8 mutations) is presented on FDC with equal frequency. Antibody feedback modulates the binding probability of antigen on FDCs. Antibodies are generated either exclusively by GC output cells (no mdPC) or, in addition, by memory derived plasma cells specific for epitope 1 (with 1k or 10k mdPC). Note that only high affinity cells to either of the epitopes are shown, i.e. the GC population is larger. Vertical bars show the standard deviation over 120 simulations for each condition.

To test whether this mechanism of self-inhibition can focus a GC reaction onto very low accessible epitope (3%), in silico GC founder cells containing 90, 25, or 5% memory B cells and variable levels of memory-derived antibody-forming plasma cells were assumed (Table 1). The more memory B cells participated in the GC, the less likely the GC responses focused on the low accessible epitope (Table 1). At first, the occurrence of B cells specific for the low accessible epitope was even lower, and secondly, those that developed were rather unlikely to be selected. According to the current belief, secondary GC reactions are dominated by memory-derived founder B cells (Pape et al., 2011). The model predicts an evolutionary advantage of individuals with a low frequency of memory B cells in GC reactions in the sense that the probability of developing bnAbs against low accessible epitopes is highest without or with only a few memory B cells initiating the GC response.

Interestingly, the presence of memory-derived antibody-forming cells specific for the 97% immunodominant epitope was able to reduce the frequency of memory B cells participating in the GC response to a similar extent as the injection of antibodies. Memory-derived antibody-forming cells increased both, the frequency of occurrence of B cell clones specific for the low accessible epitope and the probability of their selection (Table 1). A critical number of memory-derived antibody-forming cells was required to achieve the same efficiency as with antibody injections. The required strength of memory-induced self-inhibition is predicted to suppress widely participation of memory B cells in GC reactions. This might be tested in experiments in order to decide whether *in vivo* this mechanism has the potential to redirect GC responses to other epitopes. It would also be interesting to test strategies of promoting memory B cell differentiation into antibody-forming cells upon reactivation, which, according to the simulations, would strengthen antibody-feedback, suppress memory B cell survival in GC reactions, and allow the GCs to focus on so far unappreciated epitopes.

## Conclusion

It was shown before that increased numbers of Tfh cells might increase the probability of developing bnAbs (de Boer & Perelson, 2017). The present results suggest that it is possible to generate antibodies against low accessible epitopes when its relative accessibility is increased by antibodies specific for the immunodominant epitopes on FDCs, a mechanism that acts upstream of selection by Tfh cells. Thus, previous results showing that the presence of antibodies can inhibit GC reactions (Zarnitsyna et al., 2016, Zhang et al., 2013) can be turned into a positive effect if the antibodies have the right specificity. This strategy might be employed for vaccination planning on an individualized patient level. Each individual comes with his own set of pre-existing antibodies. One could complement the antibody spectrum by missing antibodies against immunodominant epitopes together with the actual vaccination and by this shift the focus of affinity maturation to hardly accessible epitopes of the conserved region. It is critical that a wide spectrum of immunodominant epitopes is covered by antibodies, such that the relative accessibility of epitopes in the conserved region is increased. The external control of specific antibody generation by injection of antibodies against immunodominant epitopes is a so far underappreciated mechanism in individualised vaccination therapy and worse being further explored.

## Acknowledgement

This work was supported by the Human Frontier Science Program (RGP0033/2015) and the Helmholtz Association, Zukunftsthema “Immunology and Inflammation” (ZT-0027). I thank Sebastian Binder for support in R-programming for generating the figures and for revising the manuscript. The author does not have any competing interests.

## Supplementary Material

### Germinal Centre simulation model

The GC model assumptions underlying the germinal centre (GC) simulations are explained here and include the used parameter values. It follows the description of (Meyer-Hermann *et al.*, 2012) adapted to include novel features introduced since then in (Meyer-Hermann, 2014; Binder & Meyer-Hermann, 2016), as well as the multi-epitope representation added here. Used acronyms are: DZ for dark zone, LZ for light zone, Tfh for T follicular helper cell, FDC for follicular dendritic cell.

### Space representation

All reactions take place on a three-dimensional discretized space with a rectangular lattice with lattice constant of Δ*x* = 5*μm*. Every lattice node can be occupied by a single cell only.

### Shape space for antibodies

Antibodies are represented on a four dimensional shape space (Perelson & Oster, 1979). The shape space is restricted to a size of 10 nodes per dimension, thus, only considering antibodies with a minimum affinity to the antigen. The optimal clone Φ* is positioned in the centre of the shape space. A position on the shape space Φ is attributed to each B cell. Its Hamming distance ||Φ – Φ*||_1_ to the optimal clone is used as a measure for the antigen binding probability. The binding probability is calculated from the Gaussian distribution with width Γ = 2.8 (Meyer-Hermann *et al.*, 2001):

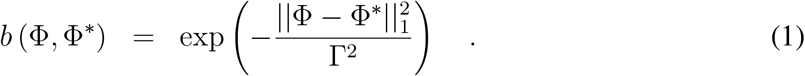

### B cell phenotypes

Three B cell phenotypes are distinguished: DZ B cells, LZ B cells, and output cells. The different phenotypes characterize the cell properties and are not meant as localization within the GC zones. DZ B cells divide, mutate and migrate. LZ B cells also migrate and undergo the different stages of the selection process. Output cells only migrate.

### Founder cells

The model starts from 250 Tfh, 200 FDCs, 300 stromal cells, and no B cell. Tfh are randomly distributed on the lattice and occupy a single node each. Stromal cells are restricted to the DZ (see section *Chemokine distribution* for their function). FDCs are restricted to the upper half of the reaction sphere, occupy one node by their soma and have 6 dendrites of 40*μm* length each. The presence of dendrites is represented as a lattice-node property and, thus, visible to B cells. The dendrites are treated as transparent for B cell or Tfh migration such that they do not inhibit cell motility.

### B cell influx rate

As B cell selection is not active during the first 3 days of the reaction (i.e. days 3-5 post immunisation), the first 3 days can be approximated as a B cell expansion phase. Clonality can be safely ignored, and it suffices to consider a single dividing cell type *B_i_*, where *i* denotes the generation of the B cells. The dynamics of expansion are then described by

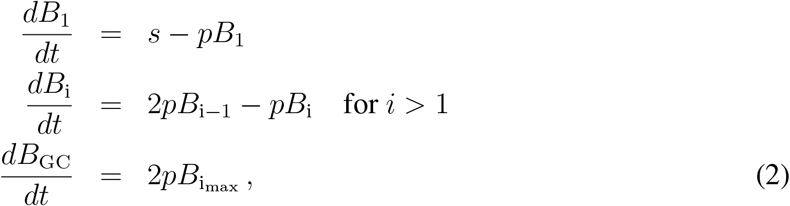

where *s* is the influx rate, *p* the division rate, *i*_max_ the number of divisions per cell in the expansion phase, and *B*_GC_ the resulting number of GC B cells that participate in the GC reaction. The number of initial divisions is estimated by the maximum number of divisions observed upon anti-DEC205-OVA treatment, i.e. *i*_max_ = 6 (Victora *et al.*, 2010; Meyer-Hermann *et al.*, 2012). As the division time ln(2)/*p* is shorter than the expansion phase *T*_expand_ = 3 days, one may solve Eq. (2) in steady state, yielding:

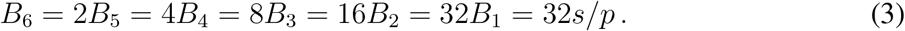

Thus, the relevant ODE becomes

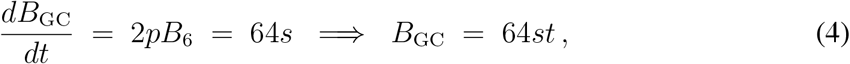

i.e. a linear growth in time proportional to the constant influx during expansion. Note that in the steady state approximation the influx rate becomes independent of the division rate *p*, which is an implication of the assumption of a fixed number of divisions per founder cell *i*_max_. With the side condition of getting 9000 cells at day 3, *B*_GC_(*T*_expand_ = 72hr) = 9000, the influx rate is estimated to be

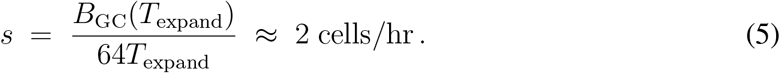

This corresponds to 144 B cells entering the GC in the first 3 days of expansion and building up the founder cell population of the GC reaction.

Motivated by this estimation, in the model we assumed that B cells enter the GC reaction with a probability corresponding to a rate of 2 cells per hour. New B cells are randomly positioned on the lattice (exclusively on free nodes).

The shape space position of each new B cell is randomly picked from a set of 100 shape space positions at a distance of 5 or 6 mutations to the nearest epitope. Memory-derived founder B cells (see below) are assumed to be optimal clones for a particular epitope, i.e. at Hamming distance zero.

### Antigen-presentation by FDCs

Each FDC is loaded with 5000 (3000 without antibody feedback) antigen portions distributed onto the lattice-nodes occupied by FDC-soma or FDC-dendrite. One antigen portion corresponds to the number of antigen molecules taken up by a B cell upon successful contact with an FDC. For multiple epitopes, both epitopes are presented at each antigen-presenting site with fractions corresponding to their accessibility.

### Antigen-antibody interaction on FDCs

The source of antibodies are either output cells of the GC reaction, memory-derived antibody-forming plasma cells, or injections (see section *Antibody sources* below). Antibodies are represented in the 4-dimensional shape space with 10 positions in each direction. The quantity of interest is the amount of free antigen-epitopes at each FDC site when antibodies are present and changing over time. As it is not feasible to calculate the amount of free epitopes at each site for all 10,000 possible antibody types, 11 affinity bins *B*_ij_ with *i* ∈ [0,…, 10] were introduced, where the affinity is defined relative to an antigen-epitope *j*, thus, every epitope gives rise to its own antibody distribution on affinity bins. We assumed a constant on-rate *k*_on_ = 10^6^/(Mol sec) (Batista & Neuberger, 1998) and a variable off-rate

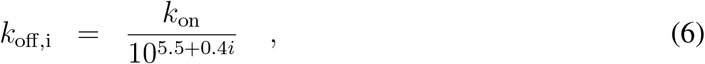

mimicking a dissociation constant that varies over 4 orders of magnitude. At each FDC site *x*, the chemical kinetics equation for the immune complexes *C*_ij_(*x*) formed between antibodies in bin *i* and epitopes *j* at the FDC site *x*

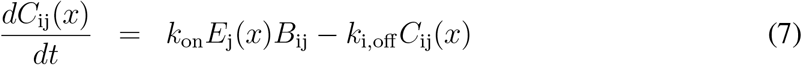

was solved for every epitope *j* in order to determine the amount of free epitope *E*_j_(*x*) at this site. Only this amount of epitope *j* is available for B cells to bind antigen with probability according to Eq. (1). No additional competitive binding probability as used in (Zhang *et al.*, 2013) was assumed here.

### DZ B cell division

The average cell cycle duration of 7 hours of DZ B cells is varied for each B cell according to a Gaussian distribution. This is needed to get desynchronization of B cell division. The cell cycle is decomposed into four phases (G1, S, G2, M) in order to localize mitotic events if this is needed.

Each founder B cell divides a number of times before differentiating to the LZ phenotype for the first time. Six divisions was the number of divisions found in response to the extreme stimulus with anti-DEC205-OVA (Victora *et al.*, 2010; Meyer-Hermann *et al.*, 2012). Each selected B cell divides an number of times determined by the interaction with Tfh (see below, LZ B cell selection). The parameters of the interaction with Tfh are tuned such that the mean number of divisions is in the range of two (Gitlin *et al.*, 2014). This value is required in order to maintain a DZ to LZ ratio in the range of two (Victora *et al.*, 2010; Meyer-Hermann *et al.*, 2012).

A division requires free space on one of the Moore neighbors of the dividing cell. Otherwise the division is postponed until a free Moore neighbor is available.

At every division the encoded antibody can mutate with a probability of 0.5 (Berek & Milstein, 1987; Nossal, 1992). This corresponds to a shift in the shape space to a von Neumann neighbor in a random direction. Upon selection by Tfh the mutation probability is individually reduced from *m*_max_ = 0.5 down to *m*_min_ = 0 in an affinity-dependent way following

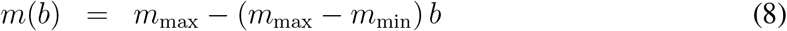

with *b* from Eq. (1) (Toellner *et al.*, 2002). Thus, after recycling DZ B cells can acquire reduced mutation probabilities. This mechanism is motivated by the observation that B cell receptor internalization enhances the activation of the kinase Akt (Chaturvedi *et al.*, 2011) which, in turn, suppresses activation induced cytosine deaminase (AID) (Omori *et al.*, 2006). AID is required for somatic hypermutation, such that this provides an affinity-dependent down-regulation of the mutation frequency (Dustin & Meyer-Hermann, 2012). However, there is no formal proof of this mechanism.

B cell division of B cells that previously acquired antigen and have been selected by Tfh distribute the retained antigen asymmetrically to the daughters (Thaunat *et al.*, 2012). The model assumes asymmetric division in 72% of the cases, which is supported by experimental observations (see (Thaunat *et al.*, 2012) and Supplementary Figure S1 in (Meyer-Hermann *et al.*, 2012)). If division is asymmetric, one daughter gets all the retained antigen while the other gets none, which approximates the value of 88% found in (Thaunat *et al.*, 2012). Mutation is suppressed in asymmetric divisions.

After the required number of divisions the B cell differentiates with a rate of in 1/6 minutes to the LZ phenotype. All B cells that kept the antigen up to this time, differentiate to output cells, up-regulate CXCR4, and leave the GC in direction of the T zone. The alternatives, that B cells randomly differentiate to output cells after divisions with a probability of 23% (LEDAX model in (Meyer-Hermann *et al.*, 2012)) or that B cells decide to differentiate to output cells right after interaction with Tfh (BASE mode in (Meyer-Hermann *et al.*, 2012)), leads to very similar GC readouts (Supplementary Figure S4). However, the amount of generated output cells is substantially higher if the B cells differentiate to output cells after divisions as compared to after selection (Dustin & Meyer-Hermann, 2012).

### LZ B cell selection

LZ B cells can be in the states *unselected, FDC-contact, FDC-selected, Tfh-contact, selected, apoptotic*.

#### Unselected

LZ B cells migrate and search for contact with FDCs loaded with antigen in order to collect antigen for 0.7 hours. If an FDC soma or dendrite is present at the position of the B cell, the B cell attempts to establish contact to the epitope with highest affinity to the B cell receptor (default setting). Alternatively, the B cell may attempt to establish contact to the epitope of highest availability at this site (used in Supplementary Fig. S3D-F). In both settings, binding is affinity dependent and happens with the probability *b* in Eq. (1). If the available number of antigen portions at the specific FDC site drops below 20 the binding probability b is linearly reduced with the number of available portions. If successful, the B cell switches to the state *FDC-contact*; otherwise the B cell continues to migrate. Further binding-attempts are prohibited for 1.2 minutes. At the end of the antigen collection period, B cells switch to the state *FDC-selected.* If a LZ B cell fails to collect any antigen at this time it switches to the state *apoptotic.*

#### FDC-contact

LZ B cells remain immobile (bound) for 3 minutes (Schwickert *et al.*, 2007) and then return to the state *unselected.* The counter for the number of successful antigen uptake events is increased by one and the FDC reduces its locally available antigen portions by one.

#### FDC-selected

B cells search for contact with Tfh. If they meet a Tfh they switch to the state *TC-contact*

#### Tfh-contact

LZ B cells remain immobile for 6 minutes. In this time the bound Tfh, which may also be bound to other B cells, polarizes to the bound B cell with highest number of successful antigen uptakes. Only this B cell receives Tfh signals and accumulate those. After the binding time, the B cell detaches and returns to the state *FDC-selected.* It continues to search and bind Tfh cells until the Tfh search time of 3 hours is over. Then, it switches to the state *apoptotic* if the accumulated Tfh-signalling time remained below 30 minutes. Otherwise it switches to the state *selected.*

#### Selected

LZ B cells keep the LZ phenotype for six hours and desensitize for CXCL13, thus, perform a random walk. During that time they re-enter cell cycle and progress through the cell cycle phases. Then they recycle back to the DZ phenotype with a rate of 1/6 minutes and memorize the amount of collected antigen as well as the cell cycle phase they have achieved by this time.

The number of divisions *P(A)* the recycled B cells will do is derived from the amount of collected antigen *A*, which reflects the amount of pMHC presented to Tfh and the affinity of the B cell receptor for the antigen, as follows:

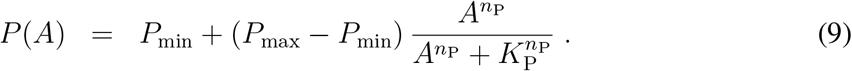

The more antigen was collected by the B cell, the more divisions are induced. We set the minimum number of division to one (*P*_min_ = 1) in order to avoid recycling events without further division. It is limited by six divisions in the best case, which is motivated by anti-DEC205-OVA experiments in which DEC205^+/+^ B cells received abundant antigen which increased pMHC presentation to a maximum (Victora *et al.*, 2010). The population dynamics in vivo and in silico only matched when the number of divisions was increased to six in the simulation (Meyer-Hermann *et al.*, 2012) suggesting that the strongest possible pMHC presentation to Tfh induces six divisions (*P*_max_ = 6). The Hill-coefficient was set to *n*_P_ = 2.

The half value *K*_P_ remained to be determined, which denotes the amount of antigen collected by B cells at which the number of divisions becomes half maximal. The number of collected antigen portions varies between zero and a maximum determined by the duration of the antigen collection phase, the duration of each B cell interaction with FDCs, and the migration time between two antigen presenting sites. The numbers of successful B cell-FDC encounters as observed in the simulations served as estimate of *A*_max_. Low affinity B cells had zero or one antigen uptake event, while high affinity cells took up between 5 and 10 portions. For an intermediate antigen uptake of *A*_0_ = 4.5, the resulting number of divisions has to be *P*_0_ = 2 in order to be in agreement with the mean number of divisions in the range of two (Gitlin *et al.*, 2014), which leads to the condition:

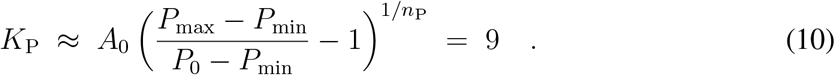

#### Apoptotic

LZ B cells remain on the lattice for 6 hours before they are deleted. They continue to be sensitive to CXCL13 during this time.

### Antibody sources

#### Output cells

Output cells from the GC reaction are recollected and memorized together with the affinity of the encoded antibody to the epitopes. Their life time is assumed longer than the duration of the GC reaction. Output cells are attributed to the different affinity bins for each epitope and further differentiate to an antibody forming plasma cell according to a linear rate equation with a rate of ln(2) per day, i.e. with a half life of one day.

All plasma cells produce antibodies *B*_ij_, attributed to bin *i* for each epitope *j* according to

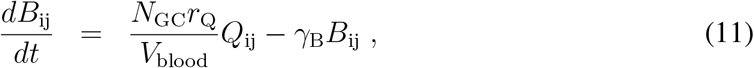

where *r*_Q_ = 3 10^−18^mol/hour is the production rate per cell (Randall *et al.*, 1992) and *γ* = ln(2)/(14days) is the degradation rate of antibodies. The produced antibodies are assumed to distribute over the whole organism. The sum of all *N_GC_* GC reactions in the organism is diluted over the mouse blood volume *V*_blood_ and adds up to the total antibody concentration found in the simulated GC. The factor *N*_GC_/*V*_blood_ = 25/ml was assumed. This setting implies that antibodies are homogeneously distributed over the space of the simulated GC. Alternative sources of antibodies are plasma cells derived from pre-existing memory B cells from a former GC reaction. Also the injection of antibodies is possible.

#### Memory-derived plasma cells

Memory-derived plasma cells are assumed to be derived from memory B cells of a previous GC response. In Figure 4, it is assumed that memory B cells specific for the immunodominant epitope pre-exist and contribute to the GC founder cells by 5% of the cells. At the beginning of the GC reaction, these memory B cells are assumed to also quickly differentiate into antibody forming plasma cells. These cells simply add to the number of antibody producing cells *Q*_ij_ in Eq. (11), i.e. *Q*_ij_ ⟶ *Q*_ij_ + *δ*(*t*)Δ*Q*_ij_, where *δ*(*t*) is the Dirac-function and Δ*Q*_ij_ = Δ*Q*_0_ for a single *i* for each epitope *j* and zero otherwise. This means that depending on the encoded antibody type, the produced antibody will be attributed to a different bin *i* for each epitope *j*. The number of memory-derived plasma cells Δ*Q*_ij_ cannot be interpreted literally, as it represents the product of a potentially different antibody production rate and the actual numbers of cells. Thus, it has to be considered as a scale for the strength of memory-derived antibody production.

#### Antibody injection

Injection of antibodies is represented by an additional source term *δ*(*t* – *t*_inject_) Δ*B*_ij_ in Eq. (11) at the particular time point *t*_inject_. Δ*B*_ij_ is Δ*B*_ij_ = Δ*B*_0_ for exactly one *i* for each epitope *j* and Δ*B*_ij_ = 0 otherwise. This means that the same amount Δ*B*_0_ is added to every epitope *j*, but to a different affinity bin *i* corresponding to the affinity of the injected antibody to each epitope.

### Chemokine distribution

Two chemokines CXCL12 and CXCL13 are considered. CXCL13 is produced by FDCs in the LZ with 10nMol per hour and FDC while CXCL12 is produced by stromal cells in the DZ with 400nMol per hour and stromal cell. As both cell types are assumed to be immobile, chemokine distributions were pre-calculated once and the resulting steady state distributions were used in all simulations.

### Chemotaxis

DZ and LZ B cells regulate their sensitivity to CXCL13 and CXCL12, respectively. This is true in all B cell states unless stated otherwise. All B cells move with a target speed of 7.5*μm/min*. This leads to a slightly lower observable average speed of ≈ 6*μm/min*.

B cells have a polarity vector that determines their preferential direction of migration. The polarity vector 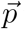 is reset every 1.5 minutes into a new direction using the chemokine distribution *c* as

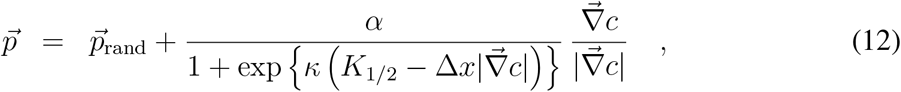

where 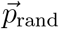 is a random polarity vector and the turning angle is sampled from the measured turning angle distribution ((Allen *et al.*, 2007) Fig. S1B). *α* =10 determines the relative weight of the chemotaxis and random walk, *K*_1/2_ = 2 · 10^11^ Mol determines the gradient of half maximum chemotaxis weight, and *κ* = 10^10^/Mol determines the steepness of the weight increase.

B cells de- and re-sensitize for their respective chemokine depending on the local chemokine concentration: The desensitization threshold is set to 6nMol and 0.08nMol for CXCL12 and CXCL13, respectively, which avoids cell clustering in the center of the zones. The resensitization threshold is set at 2/3 and 3/4 of the desensitization threshold for CXCL12 and CXCL13, respectively.

B cells can only migrate if the target node is free. If occupied and the neighbor cell is to migrate in the opposite direction (negative scalar product of the polarity vectors) both cells are exchanged with a probability of 0.5. This exchange algorithm avoids lattice artifacts leading to cell clusters.

Tfh do random walk with a preferential directionality to the LZ: The polarity vector 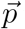 of Tfh is determined from a mixture of random walk 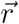 and the direction of the LZ 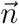 by

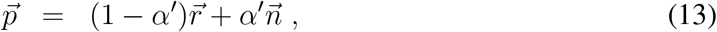

where *α’* = 0.1 is the weight of chemotaxis. This weight leads to a dominance of random walk with a tendency to accumulate in the LZ as found in experiment. TCs migrate with an average speed of 10*μm/min* and repolarize every 1.7 minutes (Miller *et al.*, 2002).

Output cell motility is derived from plasma cell motility data to be 3*μm/min* (Allen *et al.*, 2007) with a persistence time of 0.75 minutes.

---

**Supplementary Figure 1:**
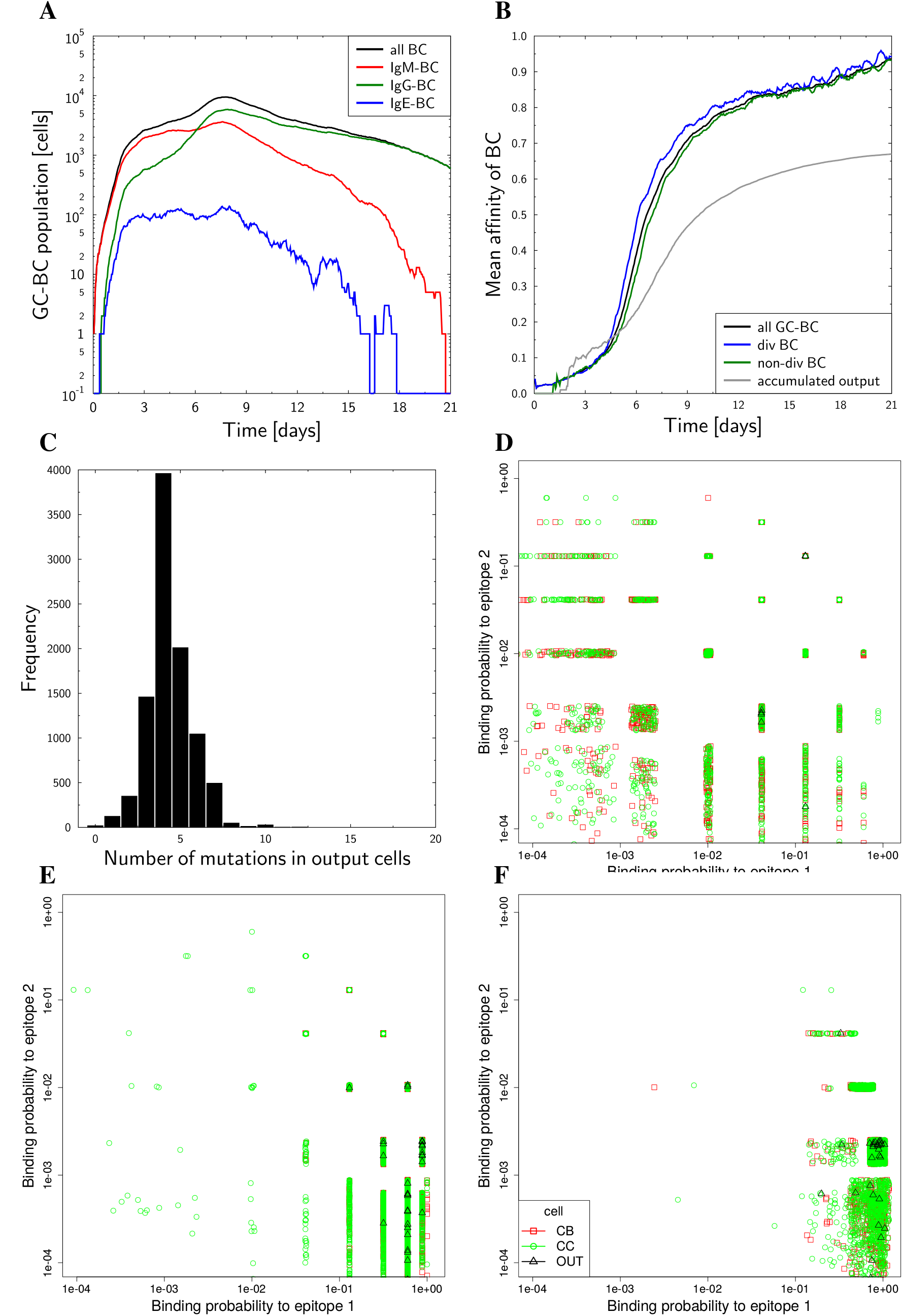
Characteristics of the reference simulation. Population kinetics (A), affinity maturation (B), the number of muations (C), and the affinity towards the presented epitope 1 on a single cell level at day 3, 7, 11, (D)-(F), respectively. Epitope 2 is not presented. Representative single simulation without antibody-feedback.

**Supplementary Figure 2:**
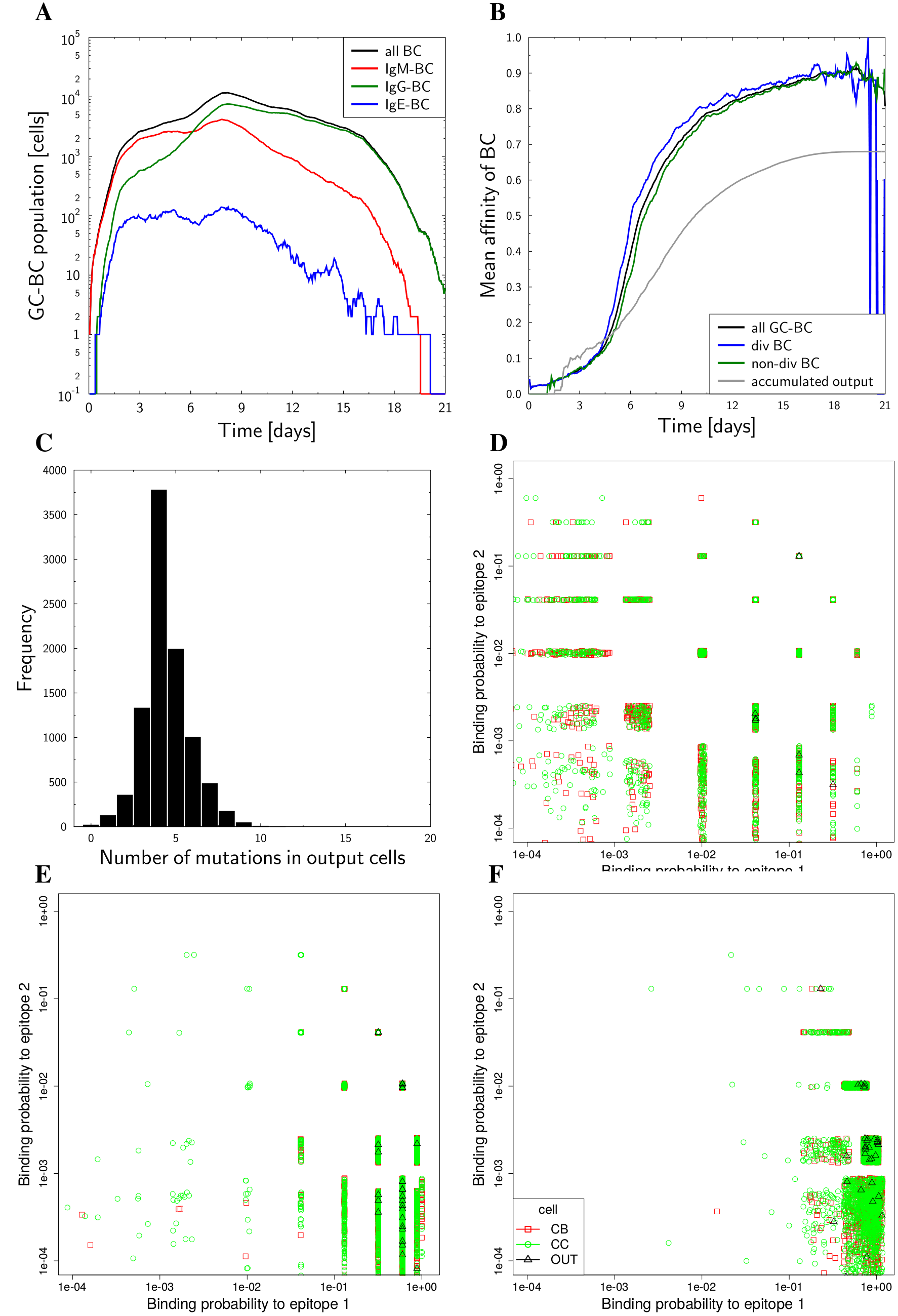
Reference simulation with antibody feedback. Population kinetics (A), affinity maturation (B), the number of muations (C), and the affinity towards the presented epitope 1 on a single cell level at day 3, 7, 11, (D)-(F), respectively. Epitope 2 is not presented. Representative single simulation with antibody-feedback.

**Supplementary Figure 3:**
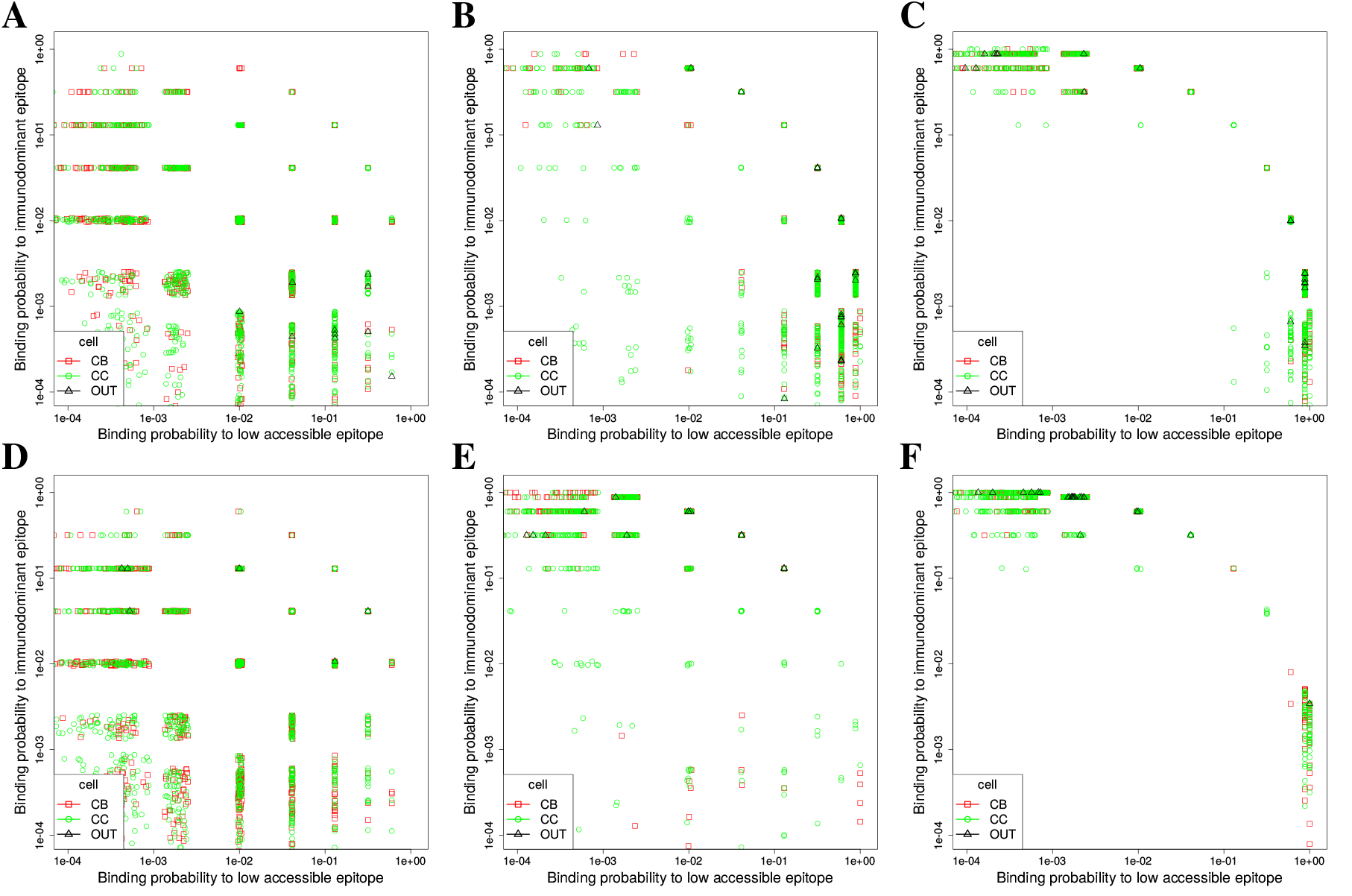
Antigen uptake models. The results in Figures 2 and 3 were generated with B cells that locally interact with the epitope of highest affinity at each FDC antigen presenting site. While this is a reasonable assumption, it was tested how the GC behaves when they interact with the epitope of highest availability at each FDC site. The evolution of B cell affinity for single representative simulations is shown at days 3, 7, and 11 of the GC reaction (in reading order). At simulation start the ratio of immunodominant and low accessible epitope is 90:10 (Hamming distance 8 mutations). B cells bind either the epitope of highest affinity (A-C) or of highest availability (D-F). 5 nMol (A-C) of antibodies specific for the immunodominant epitope is injected at day 3 of the GC reaction. For simulations (D-F), 5 nMol fully suppressed the response against the immunodominant epitope (not shown). With 2.5 nMol (see D-F) affinity maturation to both epitopes was achieved. Thus, when B cells bind the epitope of highest availability, the antibody-induced suppression of the GC response against the immunodominant epitope is even more pronounced. Less antibodies are required to achieve affinity maturation against both epitopes.

**Supplementary Figure 4:**
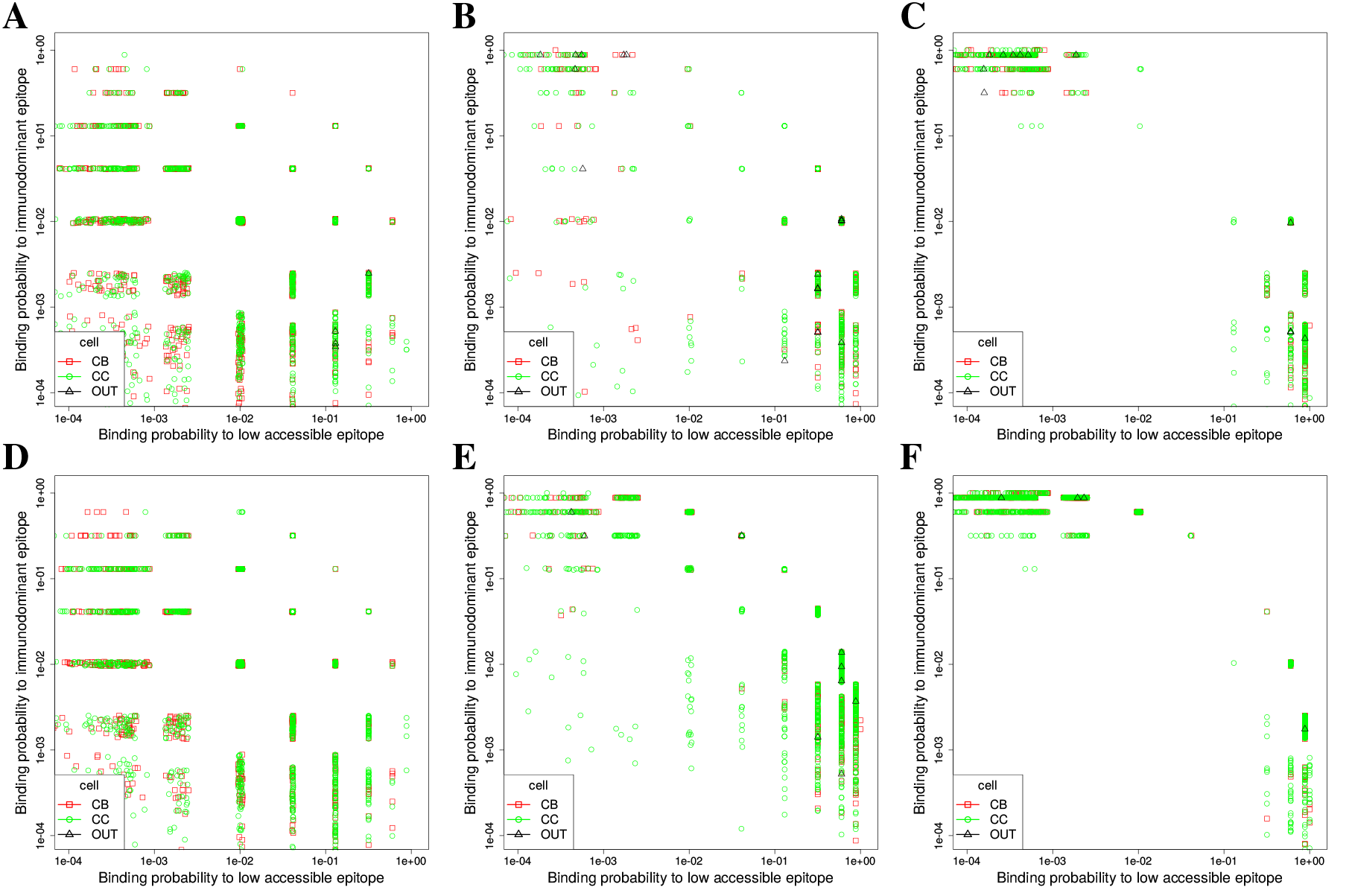
GC exit models. The results in Figures 2 and 3 were generated with the LEDA exit model (Meyer-Hermann *et al.*, 2012), according to which all selected B cells return to the dark zone, keep the collected antigen, and distribute it asymmetrically upon every division (Thaunat *et al.*, 2012). When all divisions are done, the cell which kept the antigen differentiates to output cells. Here, the antibody-injection experiment with 5nMol and with the low accessible epitope in a 10:90 ratio is repeated in two alternative models. (A-C): B cells still differentiate to output cells after divisions in the dark zone, but with a probability of 23%, irrespective of kept antigen. (D-F): 10% of the B cells differentiate to output cells right after selection in interaction with Tfh (classical exit model). In each case days 3, 7, and 11 of a single representative GC reaction are shown. Affinity maturation to the low accessible epitope is occurring in the same frequency and intensity as in the LEDA model.

